# Cerebellar Reaching Ataxia is Exacerbated by Timing Demands and Assistive Interaction Torques

**DOI:** 10.1101/2024.10.28.620711

**Authors:** Kyunggeune Oh, Di Cao, Noah J. Cowan, Amy J. Bastian

## Abstract

Individuals with cerebellar ataxia face significant challenges in controlling reaching, especially when multi-joint movements are involved. This study investigated the effects of kinematic and dynamic demands on reaching using a home-based virtual reality task. Participants with and without cerebellar ataxia reached to target locations designed to elicit a range of coordination strategies between shoulder and elbow joint movements. Compared with control subjects, cerebellar subjects presented greater initial reaching direction errors, larger hand trajectory curvatures, and more variability. Kinematic simulations indicated that early hand movement errors were sensitive to the required onset times and rates of joint movements and were most impaired when opposite direction joint movements were required (e.g., elbow extension with shoulder flexion). Dynamic analysis revealed that cerebellar participants’ movements were more impaired in reaching directions where interaction torques would normally assist the desired elbow and shoulder movements. These reach directions were also those that required joint movements in opposite directions. Overall, our data suggest that reaching deficits in cerebellar ataxia result from 1) the early-phase motion planning deficits that are exacerbated by stringent timing coordination requirements and 2) the inability to compensate for interaction torques, particularly when they assist the intended movement.

## Introduction

A hallmark of cerebellar damage is movement incoordination (ataxia). For example, people with ataxia reach with hand paths that show abnormal curvature and oscillation as they home in on a target. Many studies have quantified kinematic abnormalities in an effort to understand the fundamental motor control deficits associated with reaching ataxia. Some studies have used reductionist methods by physically constraining movement to a single-joint (e.g., elbow) and have shown kinematic abnormalities, including some dysmetria (i.e., over- or undershooting), oscillation, and prolonged deceleration phases ^1–3^. Several groups have reported that multi-jointed reaches are more impaired than single-jointed reaches, due to abnormal coordination between joints ^4–6^. Consistent with this, it has been found that a single jointed elbow movement worsens when it is made without shoulder constraint, due to poor shoulder stabilization ^7^. These findings suggest that coordination problems associated with cerebellar ataxia may stem from poor control of the more complicated kinematics and dynamics of multi-jointed movements. To examine these impairments in a broader, more diverse population, we developed and deployed a novel home-based virtual reality system that allowed participants to complete the experiment remotely from their homes.

One hypothesis is that ataxia may arise due to poor control of specific elements of movement dynamics. People with cerebellar ataxia have been reported to have difficulty predicting and accounting for “interaction torques”, torques produced at a given joint (e.g., elbow) by motions of other linked joints (e.g., shoulder) ^8–10^. Interaction torques scale with movement velocity and acceleration. They can induce large accelerations across joints, powerfully affecting reaching trajectories. Accordingly, when people with cerebellar ataxia make faster reaching movements, they show greater overshooting errors caused in part by poor compensation for interaction torques ^8–10^. Interaction torques also exhibit nonlinear characteristics, depending on the number of joints involved, their initial configurations, and relative movement directions ^7,11,12^. Importantly, depending on the reaching direction, interaction torques can assist or resist desired joint motions ^13^. Typically, adults can predictively utilize or counteract interaction torques to achieve a well-coordinated reach ^14^. It is not clear whether people with cerebellar ataxia show different patterns of deficit depending on whether interaction torques assist or resist the desired joint motions in reaching.

Impaired dynamic control may explain why people with cerebellar damage exhibit errors early in reaching that vary with movement direction ^15–18^. This has been observed in several studies that used a “center-out” reaching tasks in the horizontal plane, with equidistant targets from a central starting point, like a clock face. Using this task, people with ataxia can make early reaching errors rotated clockwise or counterclockwise relative to the desired target (e.g., Gibo et al. 2013). Each of the aforementioned studies noted that some reaching directions appeared “easier” than others—people with cerebellar damage could reach some targets directly but showed large errors for other targets. It is difficult to find a systematic pattern across these studies because they used different initial starting positions and different numbers and positions of targets and involved different robots that altered the inertia of the arm.

Here, we were interested in understanding whether people with cerebellar ataxia have different sensitivities to the kinematic and dynamic demands of reaches made with different combinations of joint movements. We used virtual reality to display reaching targets that were classified according to the direction and combination of standard 20-degree shoulder and 20-degree elbow joint motions. Some combinations require tight control of the timing and rate of shoulder and elbow kinematics for reaching, whereas others do not. In addition, some combinations involve interaction torques that assisted the desired joint motions for reaching, whereas others involve resistive interaction torques. Although interaction torques can be accentuated by fast movements, facilitating analysis ^7–10,19,20^, natural arm reaching speeds can nevertheless elicit them ^1–3^. Thus, we focus here on reaching movements that are at natural and comfortable speeds, as such movements may provide the most insight into everyday activities performed by individuals with cerebellar ataxia.

## Materials and methods

### Subjects

Seventeen control subjects and sixteen individuals with cerebellar ataxia were included in this study. All participants were right-handed, and control subjects were matched to cerebellar ataxia subjects in terms of age and sex, as detailed in Table 1. The Scale for the Assessment and Rating of Ataxia (SARA) was employed to evaluate the severity of ataxia in all cerebellar patients. The assessments were conducted remotely via Zoom video calls (Zoom Video Communications, Inc., San Jose, CA, USA), using dual-camera angles: one from the laptop’s built-in camera facing the subjects and another from a webcam positioned to their right side. Note that remote administration of the SARA has been shown to be valid and reliable ^21^. Subjects with severe ataxia were accompanied by a caregiver during the assessment. The study protocol received approval from the Johns Hopkins Medicine Institutional Review Board and was carried out in accordance with the principles outlined in the Declaration of Helsinki Prior to the study’s initiation, informed consent was obtained from all participants. A summary of the cerebellar subjects’ information can be found in Table 2.

**Table 1.** Subject information.

| **Group** | **Controls** | **Ataxia** |
| --- | --- | --- |
| Headcount | 17 | 16 |
| Age | 61.6 ($\pm 6.9$) | 60.0 ($\pm 11.9$) |
| M/F | 7/10 | 5/11 |
| Body Weight (lbs) | 164 ($\pm 26$) | 157 ($\pm 29$) |
| UA segment length (cm) | 31.7 ($\pm 2.6$) | 31.6 ($\pm 1.9$) |
| LA segment length (cm) | 39.8 ($\pm 2.8$) | 39.5 ($\pm 2.0$) |

**Table 2.**
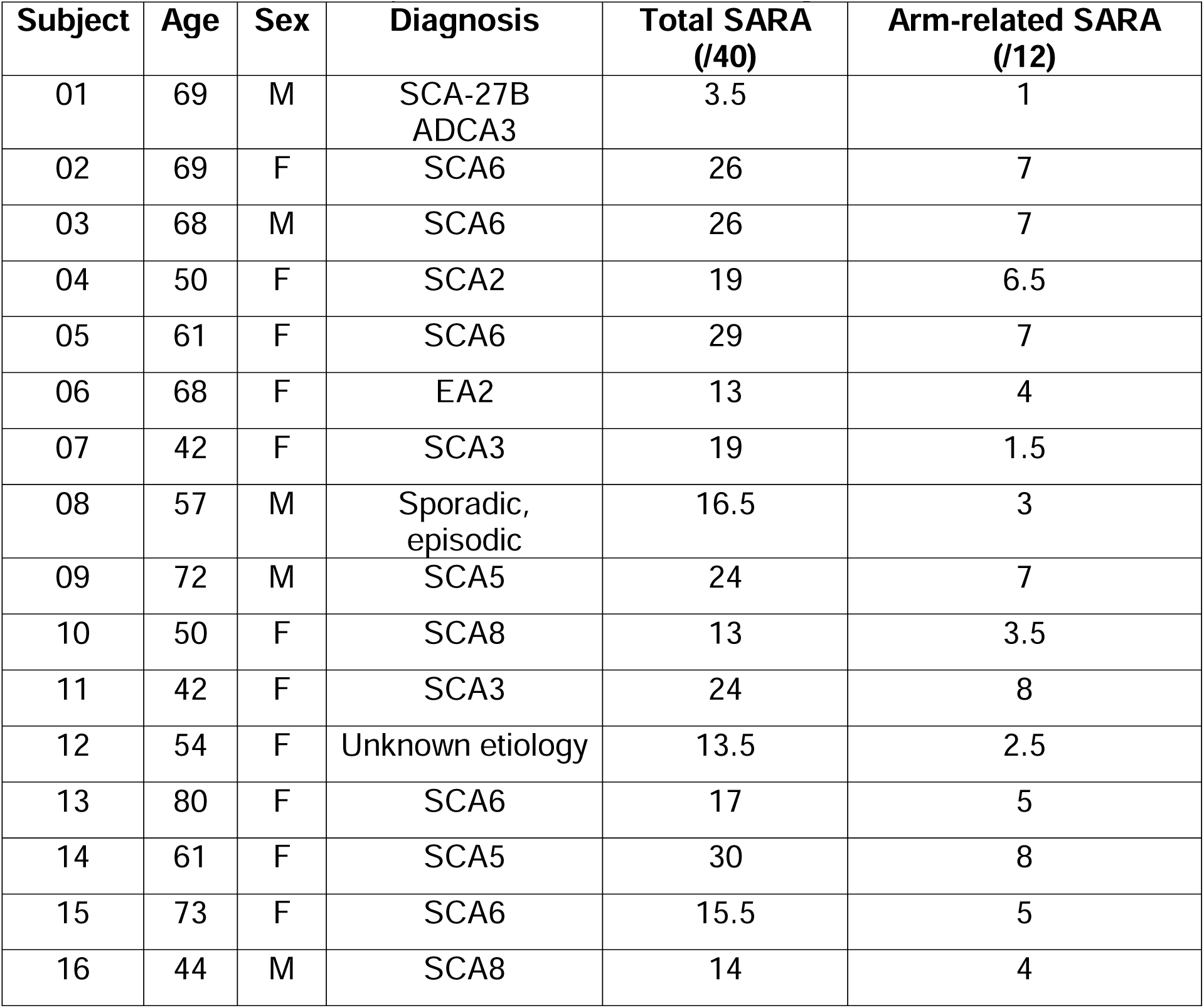
Characteristics of patients with cerebellar damage.

| **Subject** | **Age** | **Sex** | **Diagnosis** | **Total SARA (/40)** | **Arm-related SARA (/12)** |
| --- | --- | --- | --- | --- | --- |
| 01 | 69 | M | SCA-27B<br>ADCA3 | 3.5 | 1 |
| 02 | 69 | F | SCA6 | 26 | 7 |
| 03 | 68 | M | SCA6 | 26 | 7 |
| 04 | 50 | F | SCA2 | 19 | 6.5 |
| 05 | 61 | F | SCA6 | 29 | 7 |
| 06 | 68 | F | EA2 | 13 | 4 |
| 07 | 42 | F | SCA3 | 19 | 1.5 |
| 08 | 57 | M | Sporadic,<br>episodic | 16.5 | 3 |
| 09 | 72 | M | SCA5 | 24 | 7 |
| 10 | 50 | F | SCA8 | 13 | 3.5 |
| 11 | 42 | F | SCA3 | 24 | 8 |
| 12 | 54 | F | Unknown etiology | 13.5 | 2.5 |
| 13 | 80 | F | SCA6 | 17 | 5 |
| 14 | 61 | F | SCA5 | 30 | 8 |
| 15 | 73 | F | SCA6 | 15.5 | 5 |
| 16 | 44 | M | SCA8 | 14 | 4 |

### Apparatus

We designed a remote data collection system that was delivered to each participant’s home, during the COVID-19 pandemic. This allowed us to study people with cerebellar ataxia who lived far away from the lab—our subjects were located in Maryland, Virginia, Vermont, North Carolina, Tennessee, Florida, Illinois, Missouri, and California. The data collection equipment package included an Oculus Rift S virtual reality (VR) device (Meta Reality Labs, WA, USA) for hand movement measurement, a Logitech C922 webcam for monitoring subjects’ sagittal view, and a gaming laptop that was used to control the VR system, perform data collection, and to conduct the video call. The data collection rate for the webcam and VR device was 30Hz. The participants received a detailed set of instructions on how to connect all equipment with color coded cables and photographs. After the computer was connected to the internet, the remaining equipment set up was guided by investigators through a video call and remote control of the laptop using TeamViewer (Google LLC, CA, USA). During testing, the subjects sat in a stable chair with their feet on the floor wearing the Oculus Rift. The laptop sat on a stable surface in front of them and the webcam was positioned to capture a view of their right side in the sagittal plane. The hand movement was recorded by the Rift S VR, with a 30Hz sampling rate. The joint motions of the shoulder and elbow were also monitored and recorded by the webcam at a 30Hz frame rate. All ataxic participants were tested off site. The control participants were tested either off-site or onsite. If onsite, the subjects were not involved in setting up the devices. Otherwise, all testing procedures were identical.

### Subject calibration

Before beginning the reaching study, each subject was asked to have another person measure the lengths of their arm segments, including the upper arm segment (from the acromion to the radiale) and the lower arm and hand segment (from the radiale to the proximal interphalangeal joint of the index finger). This was done while we were online with the subject, providing verbal instructions and video checks. The subject was then instructed to sit on a chair and maintain a fixed trunk and head position during the task either by leaning on the back of the chair or sitting upright.

In the initial stage of the VR task, all the subjects underwent a calibration process. They put on the VR headset and held a VR controller in each hand. The participants were told that the (virtual) green arrows marked on the floor pointed in the direction that they should be facing; after verifying that they were facing the correct direction, they were asked to raise both hands (which held the VR controllers) to the height of their shoulder and extend them forward shoulder width apart. The position of the right shoulder in 3D space was estimated by recording the positions of the controllers and utilizing the measured lengths of the arm segments.

### Paradigm

The participants performed goal-directed arm reaching movements in the sagittal plane in front of their right shoulders. The starting position of the arm reach was defined as the elbow joint angle at 90 degrees and the shoulder joint angle at 40 degrees relative to the ground, as shown in Figure 1. Reaches were performed on eight targets that were displayed one at a time during the task. The targets were located in the vertical plane, with four of them being single-joint targets that could be reached with 20 degrees of excursion in either flexion or extension of the shoulder or elbow joint.

**Figure 1.**
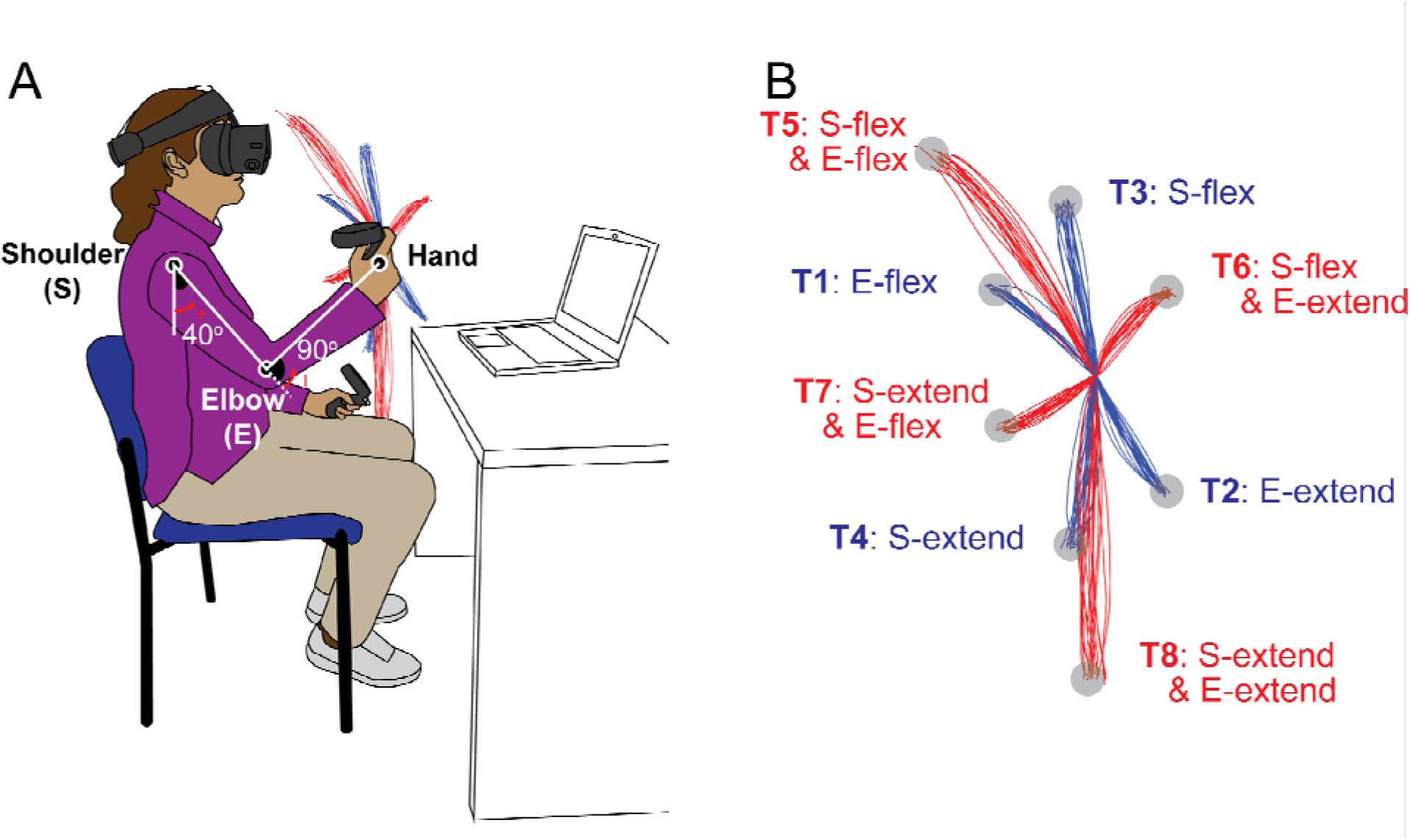
Task diagram. A. Initial position: The shoulder joint is angled at 40 degrees from the ground direction, and the elbow joint is at 90 degrees, positioning the hand at the height of the right shoulder. The positive direction for the joint angles is counterclockwise when viewed from the right. B. Target Locations: The starting point and all the targets are located on a sagittal plane that aligns with the right shoulder. The hand paths leading to four single-joint targets are illustrated in red, whereas those leading to four double-joint targets are indicated in blue.

The remaining four targets were two-joint targets, requiring 20 degrees of joint excursions in both the shoulder and elbow joints, flexion or extension. Note that we use the term “opponent joint movements” for two-joint reaches where the shoulder and elbow joints moved in opposite directions (e.g., shoulder flexion with elbow extension and vice versa) and “same-direction joint movements” when both joints were flexing or extending. A trial was defined as a single reaching movement from the starting position to one of the targets. Before data collection, the subjects performed practice trials until they became accustomed to the reaching task. For data collection, participants reached each of the 8 targets nine times, resulting in 72 total trials arranged in a pseudorandom manner.

Each trial began by displaying the starting point, a red sphere with a 2 cm radius, with a gray wire frame surrounding it. The subject’s hand (the right VR controller) position was displayed as a white sphere with a 1.5 cm radius. When the subject’s hand remained inside the red sphere for two seconds, the wire frame surrounding the starting point changed from gray to green, the target then appeared in the sagittal plane, and the starting point disappeared. At this point, the participants were instructed to reach the target with their natural arm reaching speed, using a straight hand path. All the targets were 2-cm radius spheres of different colors (other than red) and were surrounded by a wire frame whose color changed from gray to green if the hand was in the target. After two seconds, the target disappeared, and the starting point was displayed. The task was repeated in this way for a total of 72 targets. There was no time limit for each reach. The subject took breaks anytime between trials for as long as they wanted. The total task took about ∼20-30 mins, and most of the subjects took ∼1-2 breaks, which were typically over within 5 minutes.

### Data analysis

#### Hand paths

The hand position data were low-pass filtered with a 10 Hz cutoff frequency using the ‘lowpass’ function in MATLAB to reduce potential noise. Supplementary materials include example data. Velocity and acceleration were then computed from the filtered position data. To remove occasional outliers in the derived signals, a Hampel filter (MATLAB ‘hampel’ function, window size = 5) was applied. The resulting velocity and acceleration signals were then smoothed using a 5-point moving average filter via MATLAB’s ‘smooth’ function to reduce noise introduced during numerical differentiation. The onset and offset timings of each trial were defined as 10% of the peak hand velocity.

We calculated kinematic parameters within the sagittal plane that reflected different aspects of the reaching movement. The initial direction of the hand was quantified as the angle formed between a straight line from the start to the target and the hand to the target at 67 milliseconds (equivalent to 2 data frames) after the onset of the reaching movement. The initial hand direction is a critical measure as it reflects the subject’s motion planning and the efficiency of their feedforward control during the early phase of the reaching movement ^22,23^. The maximum deviation ratio was used to represent the curvature of the hand path. It was calculated as the perpendicular distance from a line connecting the start and end target to the farthest deviation of the hand path. This value was normalized to the distance from the start to the end target. A path length ratio was used to evaluate the kinematic efficiency of the hand movement. It was calculated by dividing the actual distance traveled by the hand by the straight-line distance between the start and the end target. Efficient kinematics (a lower path ratio) refers to movements that go straight to the target, and inefficient kinematics (a higher path ratio) follows a more circuitous (e.g., oscillatory) path to the target. Finally, peak hand path velocities, as well as acceleration and deceleration times, were calculated.

#### Joint angles

Joint angles were calculated using inverse kinematics assuming purely sagittal plane motion (note that the average maximum out-of-plane motion was small, 0.78 cm for the control group and 1.94 cm for the cerebellar ataxia group). The participants’ right shoulder was set as the origin of the coordinate system, with the anterior (+) – posterior (-) direction as the x-axis and the upward (+) – downward (-) direction as the y-axis. The hand position was denoted (X, Y). Using the given arm segment lengths, L1 (upper arm) and L2 (lower arm and hand), the shoulder (_sH_) and elbow (_EL_) joint angles were calculated as shown in equations (1) and (2):

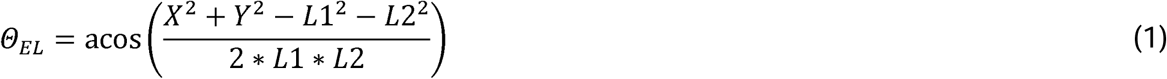

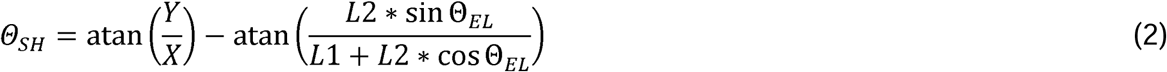

The elbow and shoulder angles are depicted in Figure 1A. Kinematic simulation A kinematic simulation was conducted to examine the impact of coordination between the shoulder and elbow joints on hand movement during two-joint reaching movements (Figure 6). First, the elbow and shoulder trajectories are time-normalized for a given target and averaged on a subject-by-subject basis. We then averaged across control subjects to create mean shoulder and elbow joint motions. These were fit to a sigmoidal curve (equation (3)) to calculate the amplitude (a), rate (r), and center point (c).

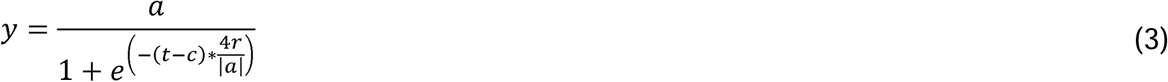

From this, we were able to systematically shift joint onset times and the relative rates of elbow and shoulder joint movements to determine the effects on hand movement deviation. Note that the hand deviation angle was calculated at the 25% point of the target distance because it provided a good estimate of the rate of shoulder and elbow motions in-flight. This was done for all combinations of simulated two-joint motions as is depicted in the heatmaps shown in Figure 6. Finally, the relative onset time difference and relative rate ratio from individual control and cerebellar subject joint motions were plotted directly on the heatmap.

Joint torque

Joint torques were calculated using inverse dynamics equations ^8^, and the calculations were based on estimations of the mass and moment of inertia for each segment, (1) the upper arm and (2) the forearm and hand ^24^. However, as wrist movement was restricted with a brace in this study, the wrist angle, angular velocity, and angular acceleration were all assumed to be zero. Alternative text: We assumed that the wrist angle, angular velocity, and angular acceleration were all zero. The inverse dynamics calculations produced the net torque (NET), gravity torque (GRAV), muscle torque (MUS), and interaction torque (INT). The basic torque equation used is NET = MUS – INT – GRAV. However, to capture the dynamic variations in the muscle torque components, we calculated the dynamic muscle torque (DMUS = MUS – GRAV), which represents the residual torque after subtracting the gravitational component ^25^. To evaluate the relationship between the dynamic muscle torque and interaction torque, we computed the zero-lag cross-correlation between the DMUS and INT.

To assess the relative contributions of DMUS and INT to NET, we calculated the contribution indices of the DMUS and INT impulses to NET ^25–27^. Note that the torque impulse represents the integral of the torque over a given time interval. The time intervals during which INT took the same sign as NET were considered to contribute a positive interaction torque impulse, whereas the intervals where the INT opposed the NET were classified as contributing to a negative interaction torque impulse. The total interaction torque impulse, obtained by summing both positive and negative impulses over the movement duration, was divided by the absolute net impulse to calculate the contribution index of the INT to the NET (equation (4)). The DMUS impulse contribution to NET was computed similarly (equation (5)).

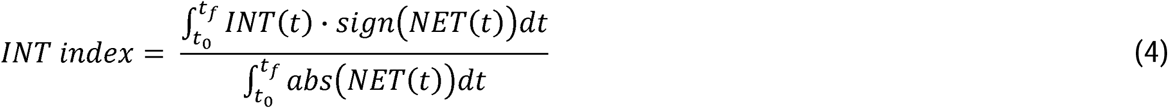

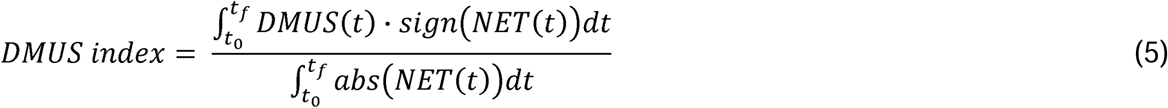

Noting that INT=NET-DMUS, the sum of the INT and DMUS indexes is always 1, as shown in equation (6):

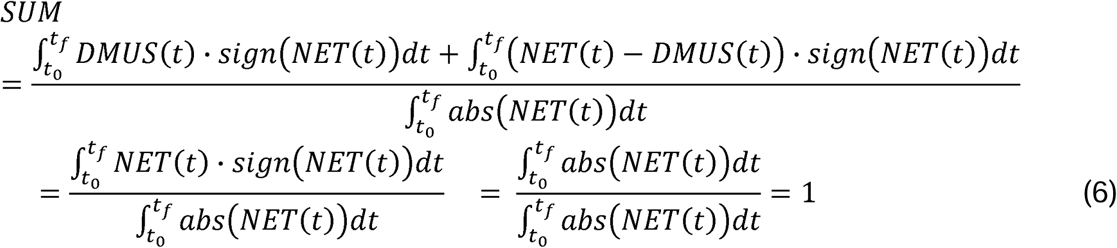

### Statistical analyses

Two-way ANOVA was conducted to examine the effects of the between-subjects factor (group: controls and cerebellar patients) and the within-subjects factor (target). Partial eta square (η^2^_P_) was reported as the measure of effect size. For post hoc comparisons, a Bonferroni correction was applied. When we examined the correlations between variables, Pearson’s correlation coefficient was employed. To ensure the robustness of this approach, we checked for homogeneity of variances between the variables using Levene’s test. All statistical analyses were performed using IBM SPSS Statistics 29 (Chicago, IL, USA).

## Results

### Hand kinematic analysis

Figure 2 shows the hand paths of the control group and the cerebellar group. A thin line traces the average hand path curve for each subject, with blue indicating paths to targets reachable via one joint movement, and red indicating those to targets requiring two joint movements. The group average curves are depicted by bold black lines. Since all the subjects had target locations adjusted for their arm segment lengths, the hand paths shown in this figure are scaled using the average target positions of each group, with the starting point positioned at the origin.

**Figure 2.**
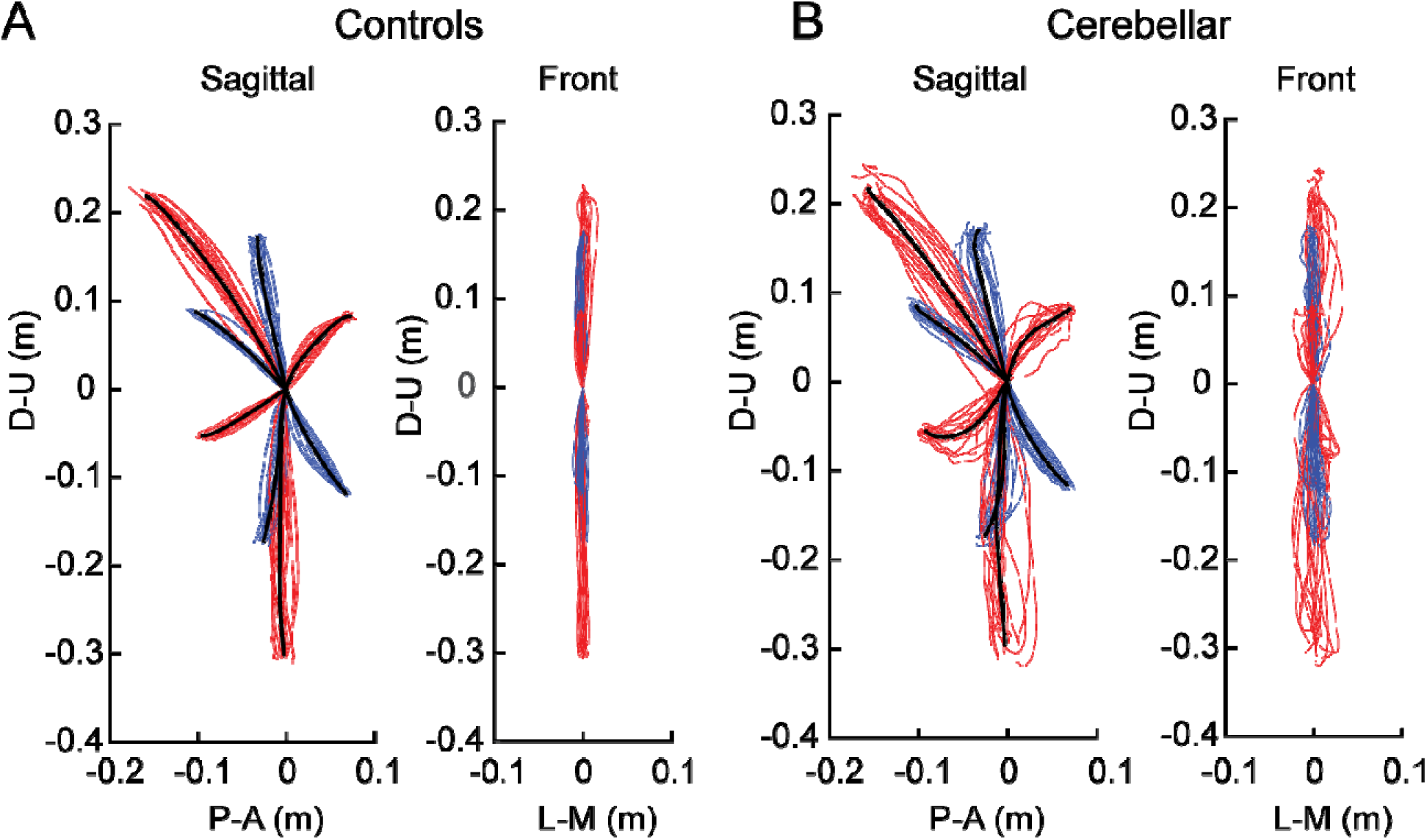
Hand paths. A. Mean hand path trajectories of all control subjects (left: sagittal view from the right; right: front view). B. Mean hand path trajectories of all cerebellar ataxia subjects (left: sagittal view from the right; right: front view). Individual lines represent the mean curves of each subject, and the thic black lines denote the mean trajectories of each subject group. The blue and red lines correspond to hand paths to single-joint and two-joint targets respectively. The hand trajectories of the different subject were scaled to the average target distances of all the subjects. Axes display directions: the Y-axis (up-down) and X-axis (anterior-posterior for the sagittal view, medial-lateral for the front view). Units are in meters.

### The cerebellar ataxia group exhibited higher inter-subject variation in hand path trajectories

In Figure 2, it is clear that the hand paths of the cerebellar group exhibited a greater degree of subject-to-subject variation than did those of the control group. This aligns with findings from previous research ^28,29^. The subject-to-subject variation in the hand paths was quantified using the standard deviation of the maximum deviation (Table 3). Additionally, hand paths viewed from the front showed greater variation in the cerebellar group; however overall, the hand paths of both groups generally remained near the target plane, as evidenced by Figure 2 (the maximum out-of-plane deviation was < 2 cm; see methods).

**Table 3.** Inter-subject variations of the hand paths: Standard deviations of maximum deviation ratio.

| Group | Target |  |  |  |  |  |  |  |
| --- | --- | --- | --- | --- | --- | --- | --- | --- |
|  | 1 | 2 | 3 | 4 | 5 | 6 | 7 | 8 |
| Control | 0.0237 | 0.0423 | 0.0224 | 0.0228 | 0.0191 | 0.0437 | 0.0223 | 0.0174 |
| Cerebellar | 0.0296 | 0.0638 | 0.0453 | 0.0612 | 0.0373 | 0.0991 | 0.1444 | 0.0362 |

The cerebellar group showed greater hand path curvature compared with the control group, and this difference was largest during opponent joint movements (targets T6 and T7).

Recall that the targets were set based on joint motions—we studied different combinations of 20-degree elbow and/or shoulder joint movements. Two of these target movements, T6 and T7, required that participants simultaneously perform flexion in one joint and extension in the other. We refer to these as *opponent joint movements* for simplicity.

Figure 2 shows that the control group’s hand paths were relatively straight for all the targets, whereas the cerebellar group’s paths were curved, a phenomenon that is consistent with similar observations reported in previous studies (Gibo et al. 2013; Zimmet et al. 2019). For both groups, the curvature was lowest in single joint movements and was greatest in two-joint movements where, one joint flexed while the other extended (and vice versa, targets T6 and T7).

In this study, the curvature of the hand path was measured using two indices, the first of which was the maximum deviation ratio, with the results displayed in Figure 3A. Generally, the cerebellar group showed significantly higher maximum deviation ratio values compared to the healthy control group (F[1, 248] = 155.993, p < 0.001, η^2^_P_=0.386), with the difference most pronounced for targets T6 and T7 (F[1, 248] = 50.062, p < 0.001, η^2^ =0.168; F[1, 248] = 145.734, p < 0.001, η^2^ =0.370). The second index utilized was the hand path ratio, which represents the relative increase in the actual path distance traveled by the hand compared with the direct distance from the starting point to the target. A perfect straight path would have a hand path ratio of 1; any deviation results in a ratio greater than 1. This metric serves as an indicator of the hand path curvature and is also interpretable as a measure of the efficiency of the reaching movement. The results, depicted in Figure 3B, demonstrate that for most targets, cerebellar patients’ hand paths were longer than those of control subjects (F[1, 248]=97.764, p < 0.001, η^2^_P_ =0.283), with the most substantial differences again observed at targets T6 and T7 (F[1, 248] = 45.091, p < 0.001, η^2^ =0.154; F[1, 248] = 48.363, p < 0.001, η^2^_P_ =0.163).

**Figure 3.**
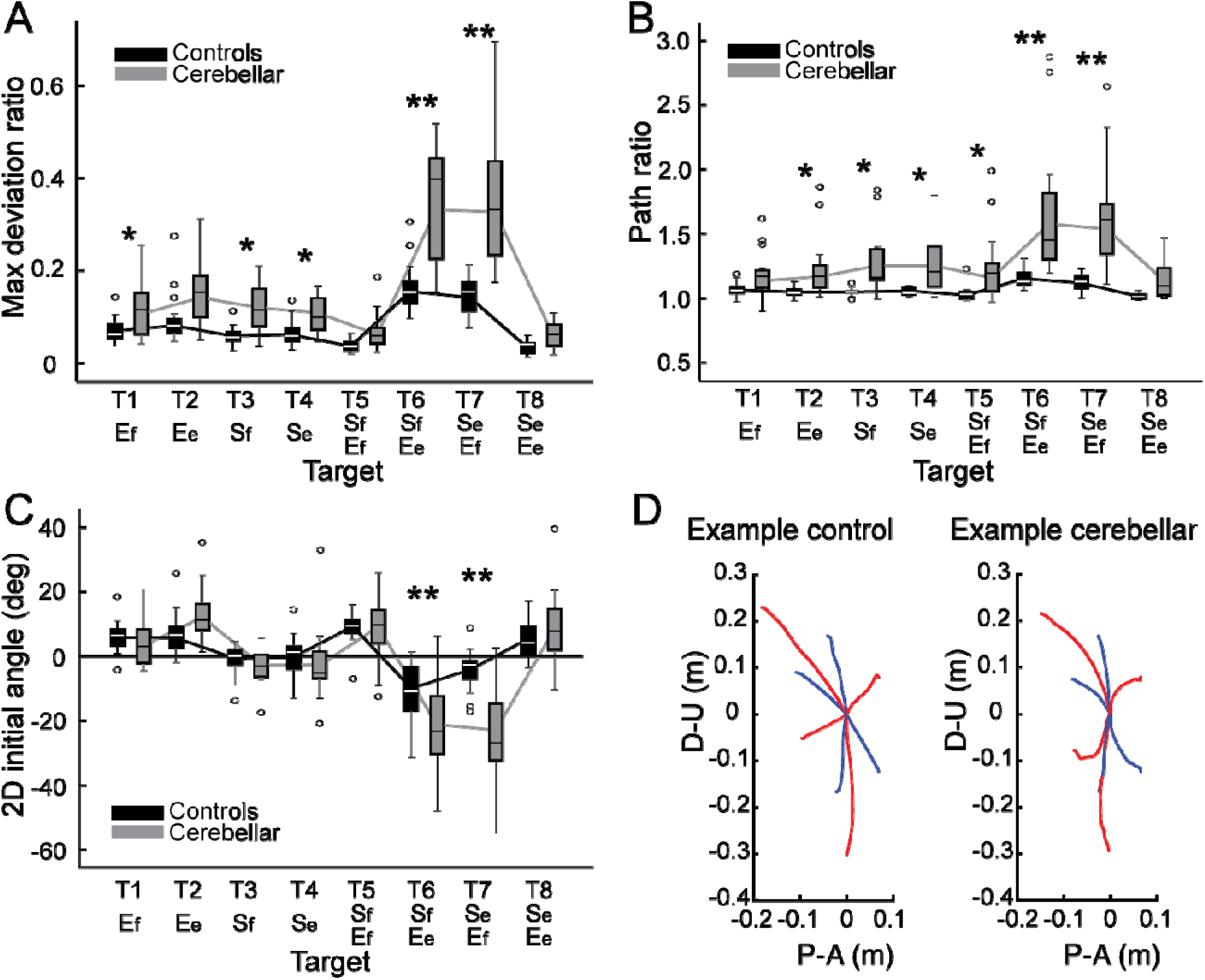
Kinematic measures of the hand path. Box plots showing the median (line), upper and lower quartiles (box), extremes (whisker) and outliers (individual points). A. Maximum deviation ratio. B. Path ratio. C. 2D initial angle. The sign of hand movement and initial direction was defined as relative to the target direction connecting the starting point and the target, with counterclockwise considered positive when viewed from the right side of the participants. *: p < 0.05, **: p < 0.001. D. Mean hand pat trajectories of representative control and cerebellar ataxia subjects. The Y-axis represents the upward (positive) and downward (negative) directions, and the X-axis represents the anterior (positive) and posterior (negative) directions, measured in meters. The blue and red lines represent hand trajectories to single-joint and two-joint targets, respectively.

### The initial hand movement misdirection was greatest for opponent joint movements (T6 and T7)

A significant difference in initial hand movement direction was observed between groups (F[1, 248] = 9.744, p = 0.002, η^2^_P_=0.038). However, this difference was particularly pronounced for targets that required coordination of opponent joint movements (T6 and T7), as shown in Figure 3C (F[1, 248] = 11.263, p < 0.001, η^2^_P_=0.043; F[1, 248] = 41.264, p < 0.001, η^2^_P_=0.143). Note that positive values are clockwise rotations, and negative values are counterclockwise rotations from a line connecting the start to target positions. Representative hand paths from the subjects in each group are illustrated in Figure 3D.

### People with cerebellar ataxia took longer to reach, with slightly different acceleration times but a markedly prolonged deceleration times

The total reach time, indicated by the entire height of the bar graph in Figure 4A, is defined as the duration from the onset to the offset of hand movement. The cerebellar group’s total reaching time was substantially longer compared with the control group (F[1, 248]=88.176, p < 0.001, η^2^_P_ =0.262), which was mainly due to a longer deceleration time.

**Figure 4.**
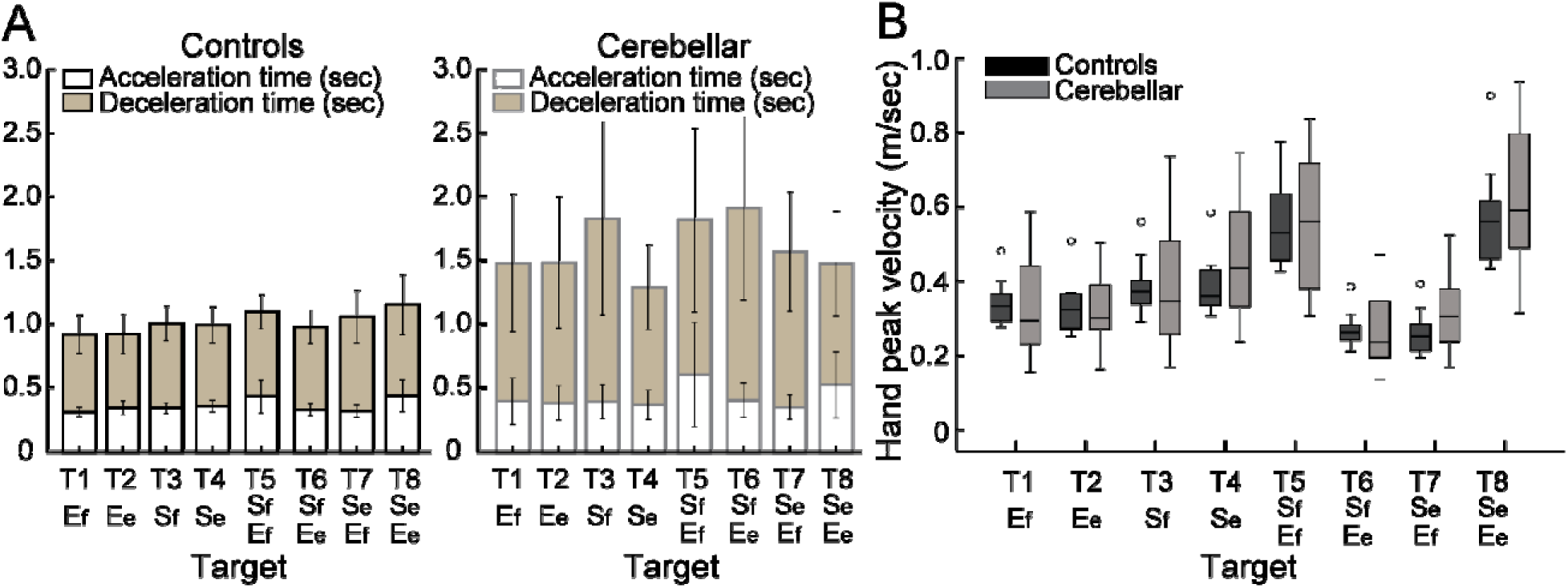
Acceleration, deceleration, and total reaching times (sec) for the control and cerebellar ataxia groups. B. Box plots for peak hand velocity (m/sec) of the controls and cerebellar groups for each target.

While the acceleration time—defined as the duration from hand movement onset to peak hand velocity—was slightly longer for the cerebellar group, 68 milliseconds, (F[1, 248]=12.717, p<0.001, η^2^_P_ =0.049), the deceleration time—defined as the period from peak hand velocity to movement offset—exhibited a significant difference of 525 milliseconds (F[1, 248]=102.903, p < 0.001, η^2^ =0.293).

### The peak hand velocities were similar between groups

As indicated in Figure 4B, the peak hand velocity between the two groups was found to be generally similar (F[1, 248]=2.955, p=0.087, η^2^_P_ =0.012). In both groups, peak velocity increased proportionally to target distance, which is consistent with findings from previous goal-directed arm reaching studies ^30^.

### Arm joint kinematics analysis

#### Joint angle trajectories showed greater inter-subject variation in the cerebellar ataxia group, and this variation was more pronounced in the elbow

Figure 5A displays the time-normalized joint angle trajectories for all the subjects. Time normalization was calculated by setting the arm reaching movement onset as 0% and the offset as 100%. Across the board, the trajectories exhibited a consistent joint excursion of approximately 20 degrees for each joint. However, in terms of trajectory variation, the cerebellar group demonstrated a greater level of variation than did the control group. When comparing joints, both groups exhibited greater joint angle trajectory variation at the elbow than at the shoulder. The variation in the joint angle trajectory was quantified using the standard deviation of the joint angle at the 50% mark on the normalized time axis, with the results depicted in Figure 5B.

**Figure 5.**
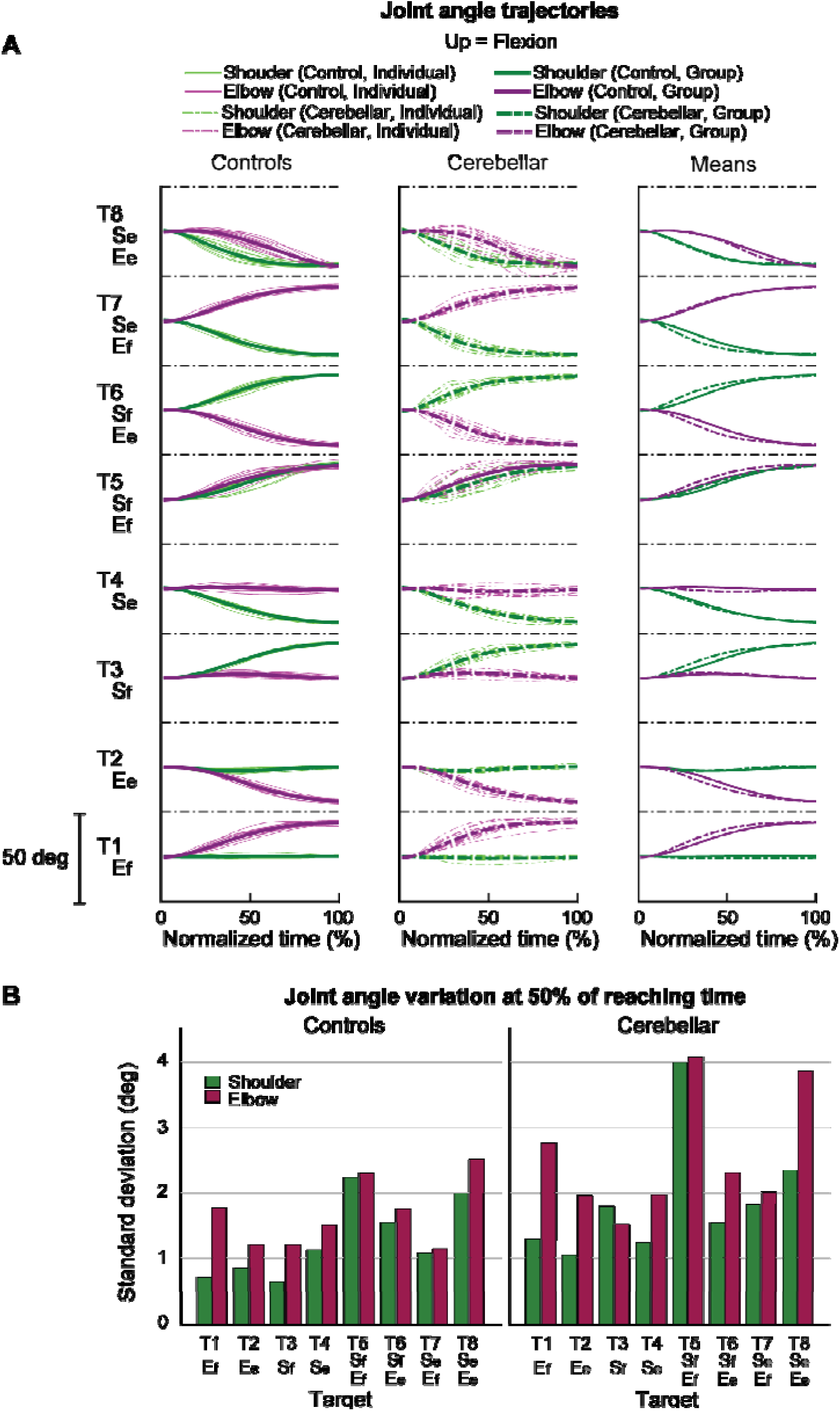
Joint angle trajectories and inter-subject variations. A. Joint angle trajectories for eight target locations, presented in the normalized time. The first and second columns display the individual mean and group mean trajectories for the control and cerebellar groups, respectively. The third column shows only group mean trajectories for both groups. B. Joint angle variation (in degrees) measured by the standard deviation of individual joint angles at 50% of the reaching time.

**Figure 6.**
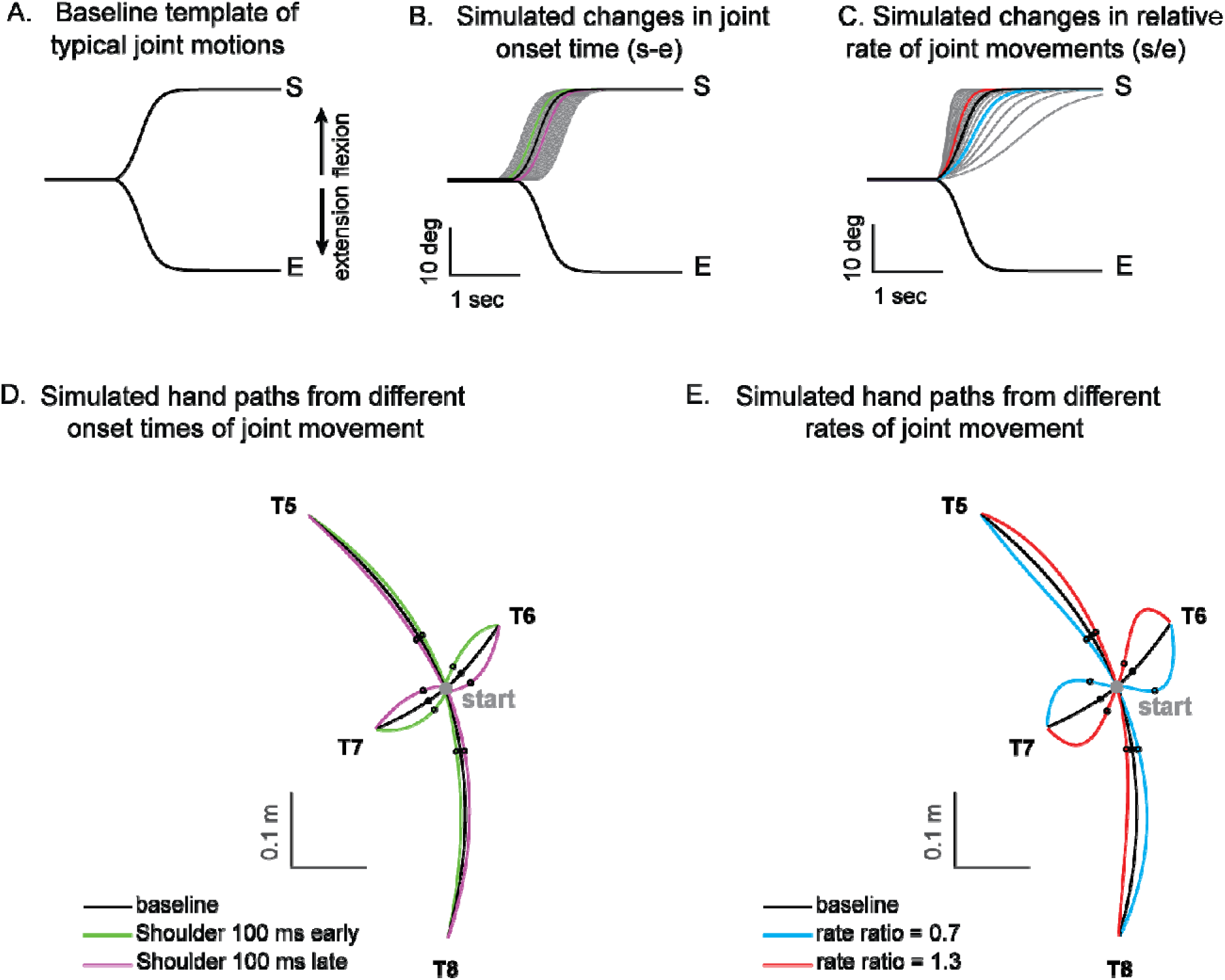
Simulated hand paths resulting from variations in shoulder and elbow joint movements. (A) Baseline template of typical joint motions (average joint motion of the control group). (B) Simulated changes in shoulder joint onset times. Shoulder joint angle trajectories with onset times 100 msec earlier or later than those of the elbow joint are shown in green and purple, respectively. (C) Simulated change in the shoulder joint angle rates. The red and blue shoulder angle trajectories represent cases where the relative rate ratios between the shoulder and elbow are 1.3 and 0.7, respectively. (D, E) Joint angle trajectories over time for each condition. The black dots indicate the points where the hand reaches 25% of the distance from the starting position in the direction of the target. At these points, the hand’s deviation angle from the target line was calculated.

### Kinematic simulations of hand deviation for two-joint targets

Kinematic simulations reveal the sensitivity of hand path deviations to different patterns of shoulder and elbow coordination. Figure 6A-C illustrates how we altered the onset times or rates of shoulder movements relative to a standard elbow movements (examples taken from T6, which required shoulder flexion and elbow extension). The joint trajectories used in this simulation were from a logistic fit to the average joint motions of the control group (see Methods). We manipulated shoulder movement to have an earlier or later onset time relative to the elbow (Figure 6B), and a faster or slower rate of movement relative to that of the elbow (Figure 6C). Figures 6D and E show example hand paths resulting from different onset times or relative rates of joint motions for all four two-joint targets (T5-T8).

Figure 7 displays the complete colormaps of the kinematic simulations. The colors in the matrix represent the degree of hand deviation from the target direction, measured in degrees, at 25% of the reaching distance. These deviations are based on variations in relative joint onset times and rate changes (slopes) between the shoulder and elbow joints in control subjects. The X-axis shows the onset time difference between the shoulder and elbow joints, and the gray vertical line represents the average onset time difference among the control subjects. The Y-axis displays the ratio of joint change rates between the shoulder and elbow joints, with the gray horizontal line representing the average rate ratio for the control group. Movements to Targets T5 and T8 were less sensitive to relative onset times and rate ratios between joints, as indicated by the gradual shift in color across the maps. In contrast, targets T6 and T7 were highly sensitive, as indicated by the abrupt changes in color across the maps.

**Figure 7.**
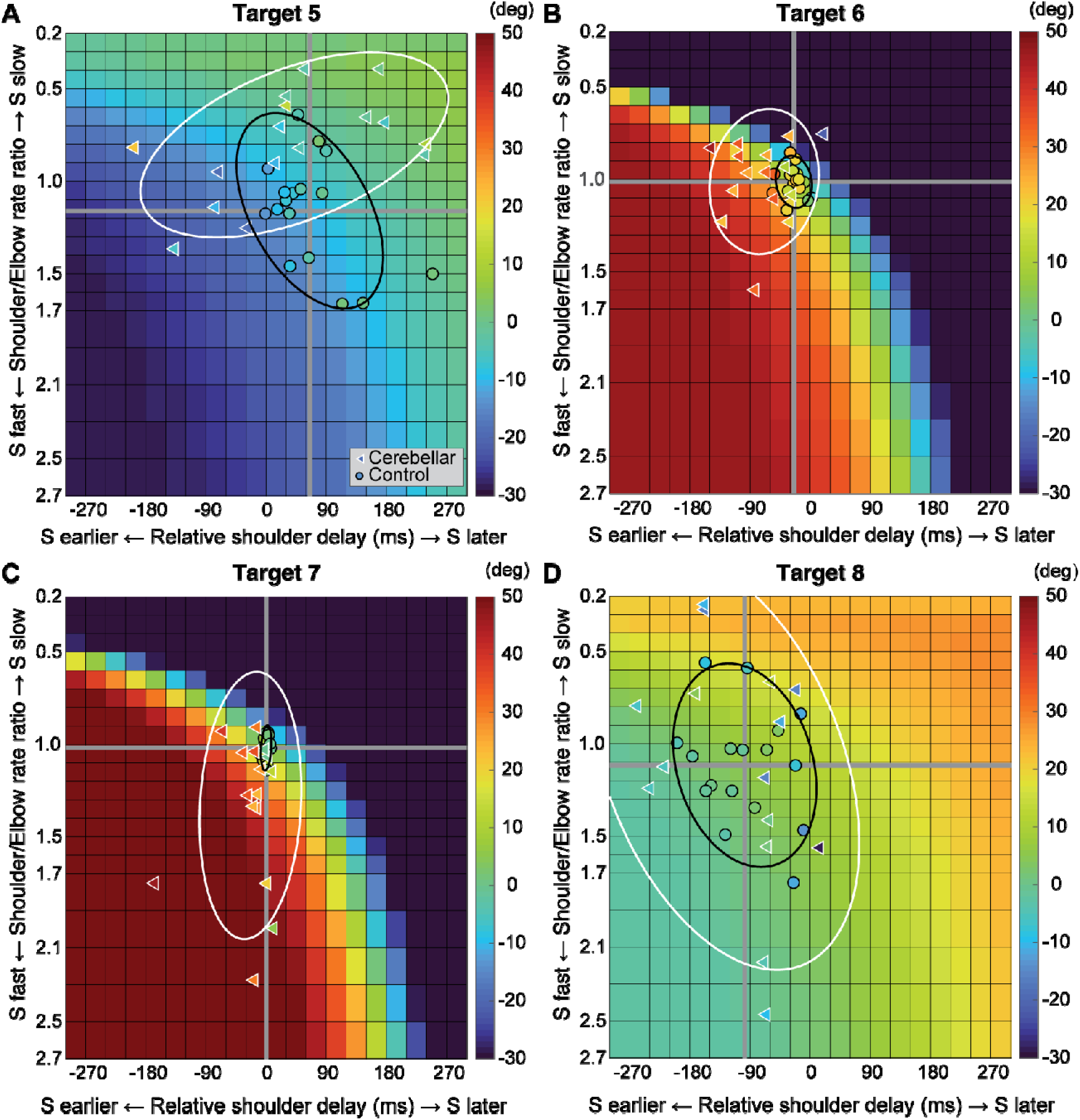
Kinematic simulations of the hand deviation from the target directions for two-joint targets: target T5 (A), T6 (B), T7 (C), and T8 (D). The simulated hand deviation from the target direction at 25% of the reaching distance is shown in degrees on colormaps and is based on variations in relative joint onset times and rate changes (slopes) between the shoulder and elbow joints in control subjects. The X-axis shows the onset time difference between the shoulder and elbow joints, where a gray vertical line represents the average onset time difference in controls. The Y-axis indicates the ratio of joint change rates between the shoulder and elbow joints, with a gray horizontal line representing the average rate ratio for the control group. Each subject’s joint onset time and joint change rate are marked individually, enabling a comparison between the predicted hand deviation from the simulation and the measured hand deviation from the experiment.

We then individually marked each subject’s joint onset time and joint change rate on the colormap, to compare the hand deviation predicted by the simulation with the hand deviation measured in the experiment. As shown in Table 4, the average difference between the two methods was 1.69 to 11.56 degrees for the control group and up to 2.62 to 15.79 degrees for the cerebellar ataxia group, depending on the target location, which can be visually confirmed by the color match between the individual data points and the colormap.

**Table 4.** Simulated vs experimentally measured hand deviation angles (deg ± standard deviation)

| Group | Target |  |  |  |
| --- | --- | --- | --- | --- |
|  | 5 | 6 | 7 | 8 |
| Healthy | -4.28 ± 4.46 | 11.56 ± 12.40 | 1.69 ± 7.55 | 10.49 ± 8.38 |
| Cerebellar | -2.62 ± 12.10 | 8.94 ± 18.32 | 5.45 ± 18.56 | 15.79 ± 13.68 |

Controls tended to show tight control of the joint onset times and rate ratios for targets T6 and T7, and looser control for targets T5 and T8. Thus, they appeared to take advantage of reduced precision requirements in joint control when they could, akin to the minimum intervention principle ^31,32^. The degree of dispersion between individual subjects’ data was represented by an 80% confidence interval error ellipse. Accordingly, for all four two-joint targets, the experimental data for the cerebellar ataxia group generally exhibited greater variation in both onset time differences and joint rate ratio differences than did the control data. For target T6, the cerebellar group showed a tendency for earlier shoulder onset times, whereas for target T7, there was a tendency for a faster relative joint angle change rate in the shoulder.

### Dynamic analysis

#### The torque patterns vary across two-jointed targets

Figure 8 presents the torque data for four two-joint targets, and illustrates the dynamic interplay between the net, interaction, and dynamic muscle torques. As shown in the figure, the arm-reaching movements performed in this study were self-paced, leading to relatively smaller interaction torques than the net torque and dynamic muscle torque. Additionally, across all four two-joint target movements, net torque and muscle torque exhibited similar patterns and magnitudes, indicating that muscle torque primarily contributed to overall movement. However, at the distal joint, the elbow, the interaction torque was often comparable in magnitude to the other torques, especially when contrasted with the shoulder. While there were no significant differences in the overall pattern between the two groups, the cerebellar ataxia group demonstrated greater inter-subject variation and less smooth torque curves

**Figure 8.**
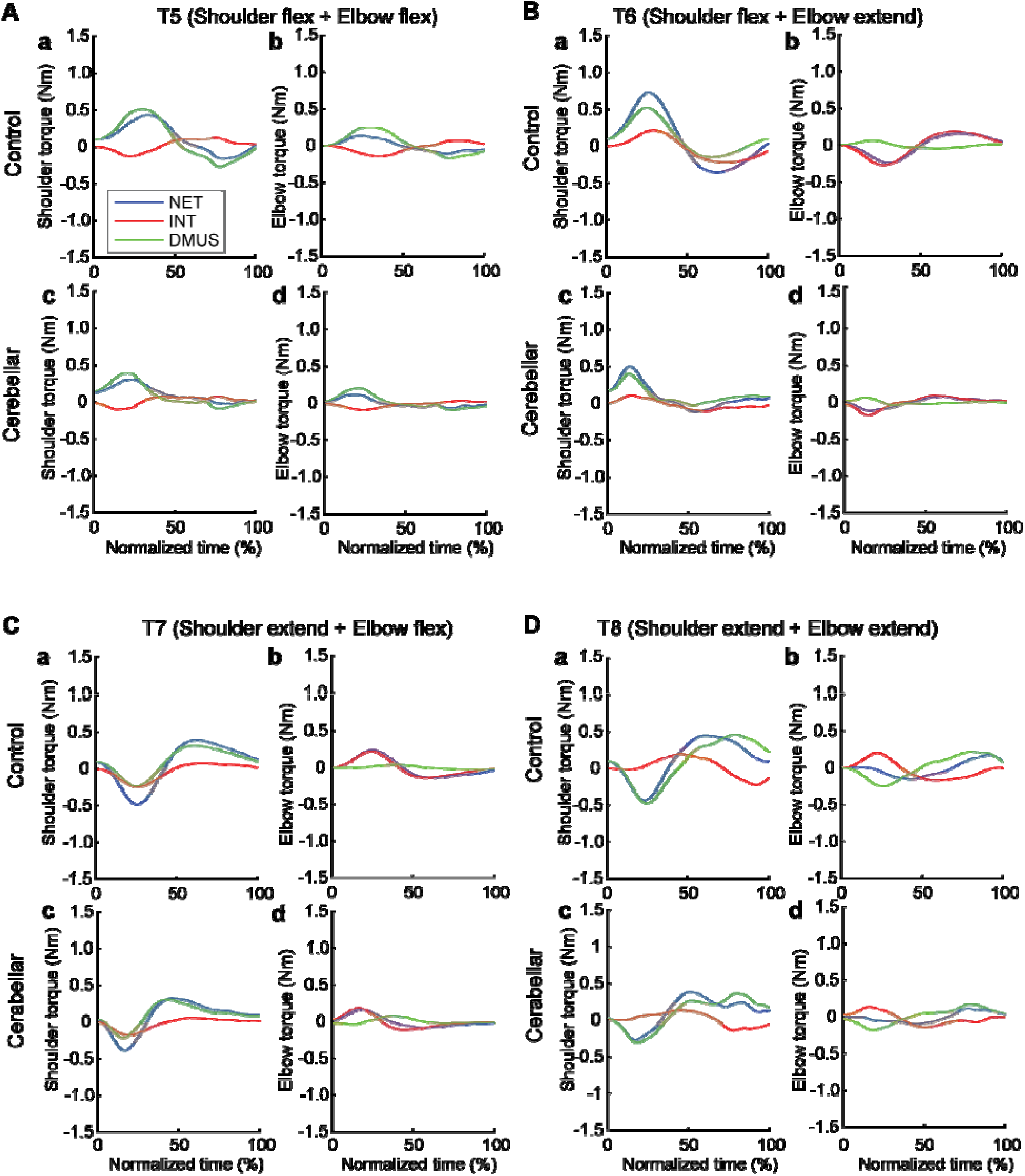
Group mean joint torque for two-joint targets: target T5 (shoulder flexion and elbow flexion, A), target T6 (shoulder flexion and elbow extension, B), target T7 (shoulder extension and elbow flexion, C), and target T8 (shoulder extension and elbow extension, D). Panel a: shoulder joint torques in the control group. Panel b: elbow joint torques in the control group. Panel c: shoulder joint torques in the cerebellar ataxia group. Panel d: elbow joint torques in the cerebellar ataxia group.

The relationship between the torques varied depending on the location of the target. Differences between the control and cerebellar groups were less apparent. For targets T6 and T7 in particular—where the shoulder and elbow joints moved in opposite directions—interaction torque in both joints appeared to positively correlate with the other torques. This would imply that for these two targets, the interaction torque acted in a manner that assisted the overall movement represented by the net torque and the movement effort shown by the muscle torque; this is examined in detail below.

#### Significant group differences were observed in the cross-correlations between interaction torque and dynamic muscle torque during opponent joint movements (T6 and T7)

To quantitatively represent the relationship between the torques, we calculated the zero-lag cross-correlation between the dynamic muscle torque and the interaction torque to examine the simultaneous relationship between the torques. As shown in Figure 9, the cross-correlation values varied depending on the target location and joint movement combination. For the shoulder joint, a positive correlation between interaction torque and dynamic muscle torque was observed for targets T6 and T7, which required opponent joint movements, suggesting that the interaction torque aided shoulder movement. In contrast, at targets T5 and T8, the interaction torque appeared to hinder shoulder movement. These findings are consistent with the torque patterns shown in Figure 8. For the elbow joint, the interaction torque generally acted in a way that hindered elbow movement. However, in the control group’s movement toward target T7, the relationship between the two showed a positive correlation, which was notably different from the cerebellar group.

**Figure 9.**
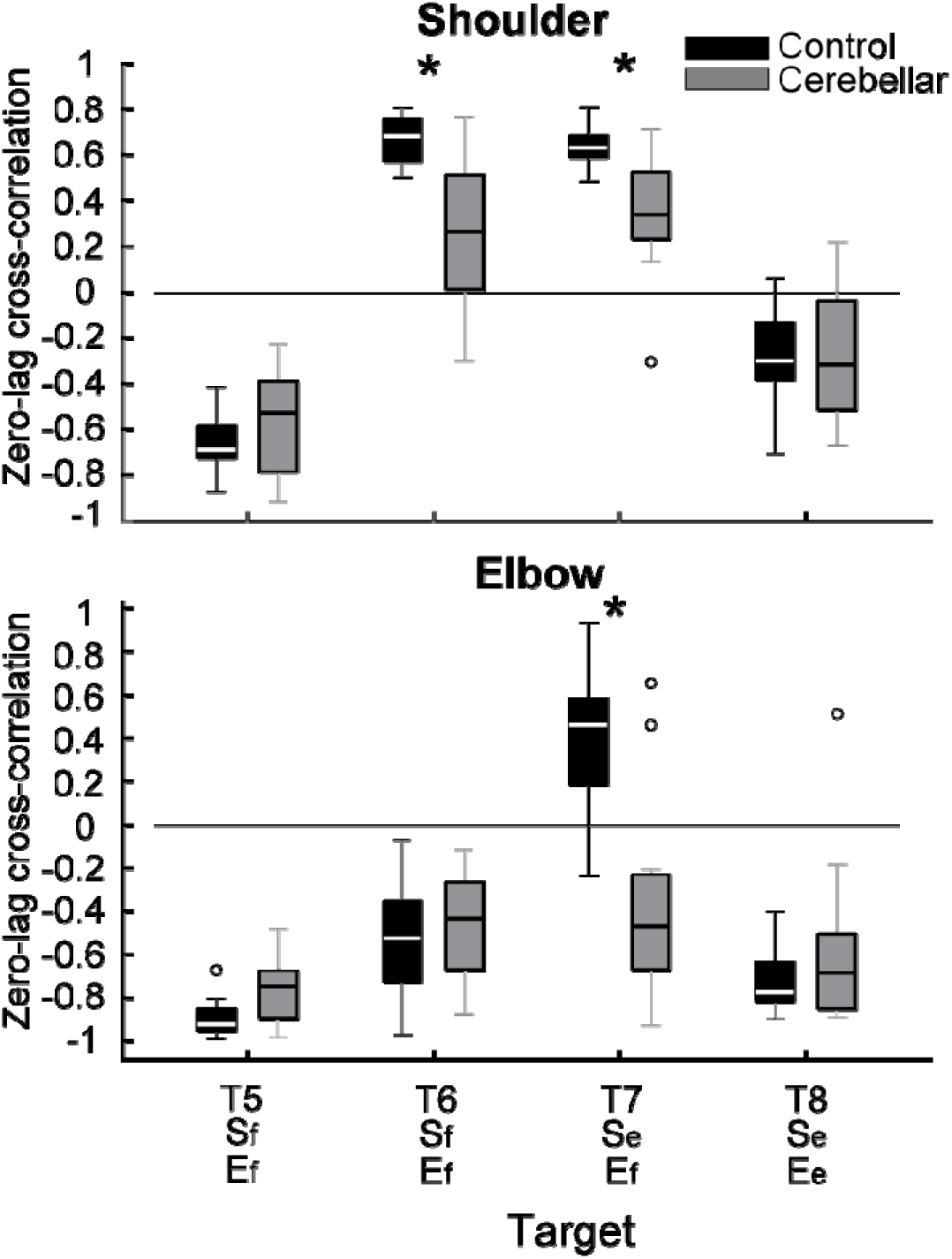
Box plots showing zero-lag cross-correlation between dynamic muscle torque and interaction torque in the shoulder and elbow joints for two-joint targets (*: p < 0.05).

The differences between the groups were most apparent at targets T6 and T7, where the shoulder joint showed a positive cross-correlation between interaction torque and dynamic muscle torque. At these two targets, the control group exhibited significantly greater cross-correlations between shoulder interaction torque and dynamic muscle torque compared with the cerebellar group (T6: F[1, 248] = 21.986, p < 0.001, η^2^_P_=0.081; T7: F[1, 248] = 11.466, p < 0.001, η^2^ =0.044). This finding indicates that the control group was able to more effectively utilize interaction torques to assist joint movement, particularly at targets requiring complex, opposite movements between the elbow and shoulder. A difference between the groups was also observed at the elbow joint for target T7 (F[1, 248] = 67.346, p < 0.001, η^2^_P_ =0.214), but this was not the case for target T6 (F[1, 248] = 0.623, p = 0.431, η^2^_P_ =0.003).

#### There was a significant reduction in interaction torque contributions during opponent joint movements made by cerebellar patients (T6 and T7)

Figure 10 shows the contribution index, which represents the relative contribution of the interaction torque and dynamic muscle torque to the net torque. Figure 8 shows that the interaction torque tended to assist the net torque for both the shoulder and elbow joints while the arm reached targets T6 and T7. This is indicated by positive contribution index values. In contrast, for targets T5 and T8, the contribution index of the interaction torque was close to zero or relatively low, indicating a minimal influence of the interaction torque on the net torque. In the case of dynamic muscle torque, both joints showed relatively high positive contribution index values, demonstrating a significant contribution to the net torque for all two-joint targets. Given the overall slower arm reaching speed, it is expected that the influence of interaction torque would be smaller than that of the dynamic muscle torque. However, during movements toward targets T6 and T7 at the elbow, the dynamic muscle torque approached zero, suggesting that interaction torque played a more dominant role in these movements.

**Figure 10.**
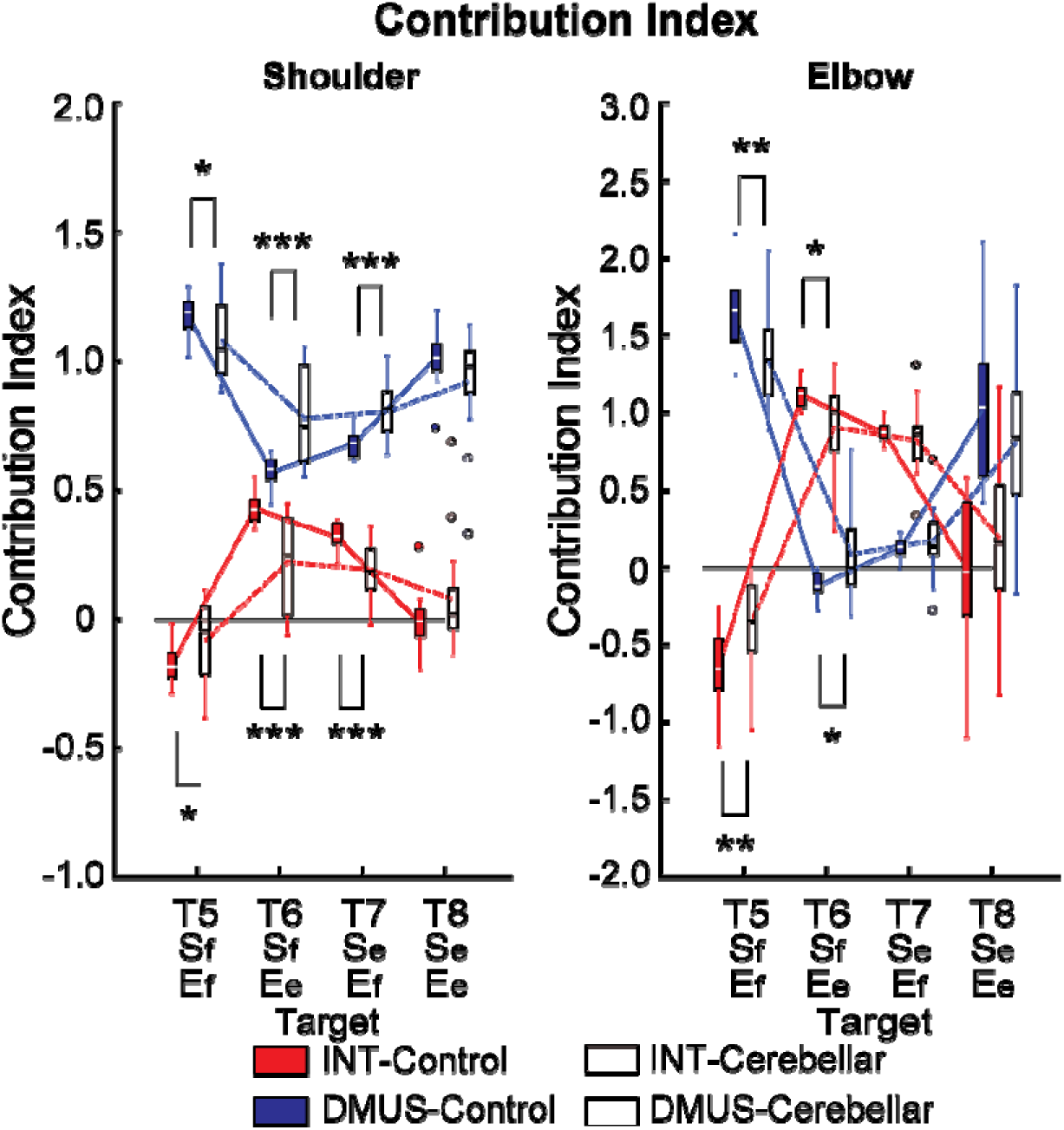
Box plots illustrating the contribution index. (*: p < 0.05, **: p < 0.01, and ***: p < 0.001).

Notably, the key difference between the two groups emerges at the shoulder joint for targets T6 and T7, where the interaction torque contributed positively to the net torque. Compared with the control group, the cerebellar ataxia group presented a significantly lower contribution index of the interaction torque to the net torque at these targets. This finding suggests that the cerebellar group had difficulty utilizing interaction torque effectively to assist movement, leading to greater reliance on dynamic muscle torque, which may not fully compensate for the movement requirements of these complex, opposite joint movement tasks. (DMUS shoulder—T5: F[1, 33] = 4.689, p = 0.038, η^2^_P_=0.131; T6: F[1, 33] = 19.346, p < 0.001, η^2^_P_ =0.384; T7: F[1, 33] = 19.229, p < 0.001, η^2^_P_=0.383; INT shoulder—T5: F[1, 33] = 4.689, p = 0.038, η^2^ =0.131; T6: F[1, 33] = 19.346, p < 0.001, η^2^_P_ =0.384; T7: F[1, 33] = 19.229, p < 0.001, η^2^ _P_ =0.383; DMUS elbow—T5: F[1, 33] = 8.585, p = 0.006, η^2^_P_ =0.217; T6: F[1, 33] = 7.108, p < 0.012, η^2^_P_=0.187; INT elbow—T5: F[1, 33] = 8.585, p = 0.006, η^2^_P_ =0.217; T6: F[1, 33] = 7.108, p < 0.012, η^2^_P_ =0.187).

## Discussion

In this study, we investigated whether people with cerebellar damage exhibit specific sensitivities to kinematic and dynamic demands during reaching. We examined participants with cerebellar ataxia and matched controls as they performed reaching task in virtual reality, using either single-joint or two-joint arm movements. As expected, controls made near-straight hand trajectories to all the targets, with low inter-subject variability. In contrast, cerebellar subjects made curved hand trajectories that varied across targets, with high inter-subject variability.

We were surprised to find that the cerebellar group had more impairments in the two-joint reaches that involved opponent joint movements than in movements where both joints moved in the same direction. All two-joint reaches involve coordination between joints and generated interaction torques, so we expected them to be comparably impaired. Instead, we found that opponent two-joint reaches were more sensitive to kinematic timing demands than same-direction two-joint movements were. Our kinematic simulation supports this—a similar error in joint onset times and rates of motion would manifest differently depending on whether the joint movements occur in the opposite or the same direction. For example, poor timing of opponent joint movements would not move the hand closer to the target and may even move the hand away from it. The same-direction joint movements tend to move the hand approximately toward the target, so inter-joint timing matters less. Single joint reaches were also less impaired in the cerebellar group, regardless of the joint used (elbow or shoulder) or the direction of movement (flexion or extension).

We also found that the dynamics of the two-joint reaches were different. In the opponent two-joint reaches, the interaction torques assisted the movement—they were positively correlated with the dynamic muscle torques and worked together to generate the movement. The cerebellar group had difficulty taking advantage of the assistive interaction torques. Compared with the controls, the cerebellar group showed lower correlations between interaction and dynamic muscle torques with smaller contributions of the interaction torques to overall joint movements. We speculate that assistive interaction torques would be difficult to incorporate into a movement if an individual has difficulty predicting how they will move the arm, possibly due to misattribution of joint-specific forces, as described by Singh and Scott (2003), who showed that under uncertainty, loads are often incorrectly associated with the motion of the joint where they are felt ^33^. The cerebellar group was better able to counter resistive interaction torques during same-direction two-joint reaches and during single jointed movements to stabilize the stationary joint. Resistive interaction torques may be easier for our cerebellar subjects to counter through feedback control in response to the perturbations that they cause ^1^.

Note that these deficits were not clearly related to the velocity of the reaching movements. It is well known that cerebellar subjects show greater deficits with faster movements ^8,9^. Here, the peak reach velocities varied depending on the reaching movement. This is because the hand-to-target distance varied with different combinations of 20° joint motions. However, the most impaired (i.e., opponent two-joint) reaches were shorter and thus hit lower peak velocities. Overall, we found that the peak hand velocities and acceleration times were slightly different between the control and cerebellar groups, suggesting that speed was not the prime factor explaining our results. We did observe longer hand deceleration times in the cerebellar group than in the control group, which has been previously reported, particularly when end-point target accuracy is prioritized ^8,34^

It is interesting that the reaches which were most difficult for our cerebellar group exhibited both features that we identified as problematic: greater kinematic temporal demands and assistive interaction torques. We think that both of these factors were important challenges for the cerebellar group, but our study does not allow us to determine which one is most important. We hypothesize that either would make reaching harder to control. Future studies should design reaching tasks specifically to disentangle these factors more clearly. We also expect that the cerebellar participants may have greater difficulty controlling the interaction torques in faster movements where the dynamics demands become much more salient.

In addition, we acknowledge that our cerebellar ataxia group was clinically heterogeneous, including individuals with subtypes such as SCA3. We examined these individuals on a video call and looked for overt signs of extra-cerebellar dysfunction, but we cannot rule out damage to other systems. As such, the observed impairments may not reflect pure cerebellar dysfunction alone. This heterogeneity is a limitation of the current study and may affect the generalizability of our findings.

Home-based studies such as these inevitably have several limitations. We reported that the cerebellar group showed out-of-plane movements that were small, but still larger than those that we observed in the control group. We acknowledge that this could introduce very small errors in our inverse kinematics calculations of sagittal plane joint motions, but did not affect our overall results. For example, a person who had the average arm length of our study participants (upper arm = 30 cm, lower arm and hand = 40 cm) in a posture with the elbow at 90° and the shoulder at 45° would have < 0.1° error at both joints if the hand moved 3 cm out of the sagittal plane. Another limitation is that the kinematic simulations were not perfectly matched with the human data. This is because our simulations used idealized joint movements that were based on logistic fits of the average control group reaches. This simplification captures general patterns but does not account for the idiosyncrasies of how individual subjects move (e.g., changes in the rate of joint motions during the reach). We believe that this method still provides a picture of the overall sensitivities of the different two-joint reaches. Finally, while the sampling rate of the oculus rift was lower than what we would typically use in the laboratory, it was more than sufficient to capture the self-paced reaches studied here. Although it is self-evident that higher sampling frequencies generally provide better data quality, our results suggest that the 30 Hz rate was adequate for capturing key movement features in this context. This was further supported by our data processing approach, which involved low-pass filtering (cutoff: 10 Hz) and additional smoothing steps to extract reliable velocity and acceleration signals from position data. Representative examples of raw and filtered data are provided in the Supplementary Materials.

In summary, the kinematic and dynamic results suggest that cerebellar subjects’ reaching ataxia stems not only from impaired timing of joint motions but also from their inability to effectively harness assistive torques that act across joints. This inability to properly coordinate both kinematics and dynamics results in the observed pattern of reach trajectories. Thus, rehabilitation strategies that focus on enhancing both the timing of joint movements and the ability to manage interaction torques could be particularly beneficial for people with cerebellar ataxia.

## Data availability

The datasets generated and analyzed during the current study are available from the corresponding author upon reasonable request. Controlled access to data is required to protect participant privacy.

## Author contributions

Conceptualization: K.O., D.C., N.J.C., and A.J.B.; Data collection: K.O.; Data analysis: K.O.; Writing-original: K.O.; Writing-review and editing: K.O., D.C., N.J.C., and A.J.B.

## Competing interests

The authors declare no competing interests.

## Funding

This work was supported by the National Institute of Health R01 HD040289 to A. J. Bastian.

## Figure legends

Figure 11. Task diagram. A. Initial position: The shoulder joint is angled at 40 degrees from the ground direction, and the elbow joint is at 90 degrees, positioning the hand at the height of the right shoulder. The positive direction for the joint angles is counterclockwise when viewed from the right. B. Target Locations: The starting point and all the targets are located on a sagittal plane that aligns with the right shoulder. The hand paths leading to four single-joint targets are illustrated in red, whereas those leading to four double-joint targets are indicated in blue.

Figure 12. Hand paths. A. Mean hand path trajectories of all control subjects (left: sagittal view from the right; right: front view). B. Mean hand path trajectories of all cerebellar ataxia subjects (left: sagittal view from the right; right: front view). Individual lines represent the mean curves of each subject, and the thick black lines denote the mean trajectories of each subject group. The blue and red lines correspond to hand paths to single-joint and two-joint targets respectively. The hand trajectories of the different subjects were scaled to the average target distances of all the subjects. Axes display directions: the Y-axis (up-down) and X-axis (anterior-posterior for the sagittal view, medial-lateral for the front view). Units are in meters.

*Figure 13. Kinematic measures of the hand path. Box plots showing the median (line), upper and lower quartiles (box), extremes (whisker) and outliers (individual points). A. Maximum deviation ratio. B. Path ratio. C. 2D initial angle. The sign of hand movement and initial direction was defined as relative to the target direction connecting the starting point and the target, with counterclockwise considered positive when viewed from the right side of the participants. *: p < 0.05, **: p < 0.001. D. Mean hand path trajectories of representative control and cerebellar ataxia subjects. The Y-axis represents the upward (positive) and downward (negative) directions, and the X-axis represents the anterior (positive) and posterior (negative) directions, measured in meters. The blue and red lines represent hand trajectories to single-joint and two-joint targets, respectively.*

*Figure 14. Acceleration, deceleration, and total reaching times (sec) for the control and cerebellar ataxia groups. B. Box plots for peak hand velocity (m/sec) of the controls and cerebellar groups for each target.*

Figure 15. Joint angle trajectories and inter-subject variations. A. Joint angle trajectories for eight target locations, presented in the normalized time. The first and second columns display the individual mean and group mean trajectories for the control and cerebellar groups, respectively. The third column shows only group mean trajectories for both groups. B. Joint angle variation (in degrees) measured by the standard deviation of individual joint angles at 50% of the reaching time.

Figure 16. Simulated hand paths resulting from variations in shoulder and elbow joint movements. (A) Baseline template of typical joint motions (average joint motion of the control group). (B) Simulated changes in shoulder joint onset times. Shoulder joint angle trajectories with onset times 100 msec earlier or later than those of the elbow joint are shown in green and purple, respectively. (C) Simulated changes in the shoulder joint angle rates. The red and blue shoulder angle trajectories represent cases where the relative rate ratios between the shoulder and elbow are 1.3 and 0.7, respectively. (D, E) Joint angle trajectories over time for each condition. The black dots indicate the points where the hand reaches 25% of the distance from the starting position in the direction of the target. At these points, the hand’s deviation angle from the target line was calculated.

Figure 17. Kinematic simulations of the hand deviation from the target directions for two-joint targets: target T5 (A), T6 (B), T7 (C), and T8 (D). The simulated hand deviation from the target direction at 25% of the reaching distance is shown in degrees on colormaps and is based on variations in relative joint onset times and rate changes (slopes) between the shoulder and elbow joints in control subjects. The X-axis shows the onset time difference between the shoulder and elbow joints, where a gray vertical line represents the average onset time difference in controls. The Y-axis indicates the ratio of joint change rates between the shoulder and elbow joints, with a gray horizontal line representing the average rate ratio for the control group. Each subject’s joint onset time and joint change rate are marked individually, enabling a comparison between the predicted hand deviation from the simulation and the measured hand deviation from the experiment.

Figure 18. Group mean joint torque for two-joint targets: target T5 (shoulder flexion and elbow flexion, A), target T6 (shoulder flexion and elbow extension, B), target T7 (shoulder extension and elbow flexion, C), and target T8 (shoulder extension and elbow extension, D). Panel a: shoulder joint torques in the control group. Panel b: elbow joint torques in the control group. Panel c: shoulder joint torques in the cerebellar ataxia group. Panel d: elbow joint torques in the cerebellar ataxia group.

*Figure 19. Box plots showing zero-lag cross-correlation between dynamic muscle torque and interaction torque in the shoulder and elbow joints for two-joint targets (*: p < 0.05).*

*Figure 20. Box plots illustrating the contribution index. (*: p < 0.05, **: p < 0.01, and ***: p < 0.001).*

## Notes

### Competing Interest Statement

The authors have declared no competing interest.

### Summary of Updates

Data Processing Clarification: We clarified that no interpolation was applied to the 30 Hz hand position data due to the slow movement speed. Instead, the data were processed using a low-pass filter (10 Hz), Hampel filtering, and moving average smoothing. Representative raw and filtered waveforms have been added to the Supplementary Materials. Effect Sizes Added: Partial eta-squared values were included for all relevant ANOVA results, and details were added to the Statistical Analyses section. Group Heterogeneity Acknowledged: We noted the heterogeneity within the cerebellar ataxia group and its implications in the Discussion. Improved Data Visualization: Figures 3, 4B, 9, and 10 were revised to include box plots with individual data points to better reflect participant-level variability. Title Revised: The title was updated to: Cerebellar Reaching Ataxia is Exacerbated by Timing Demands and Assistive Interaction Torques. Relevant Literature Cited: Singh & Scott (2003) was cited to support discussion of torque misattribution in uncertain conditions. Home-Based Data Collection Emphasized: The abstract and introduction were revised to highlight the use of a novel home-based virtual reality system. Minor Edits and Formatting: All identified typos, duplicated text, and formatting inconsistencies were corrected throughout the manuscript.

