## Supplementary materials for "Cerebellar Reaching Ataxia is Exacerbated by Timing Demands and Assistive Interaction Torques"

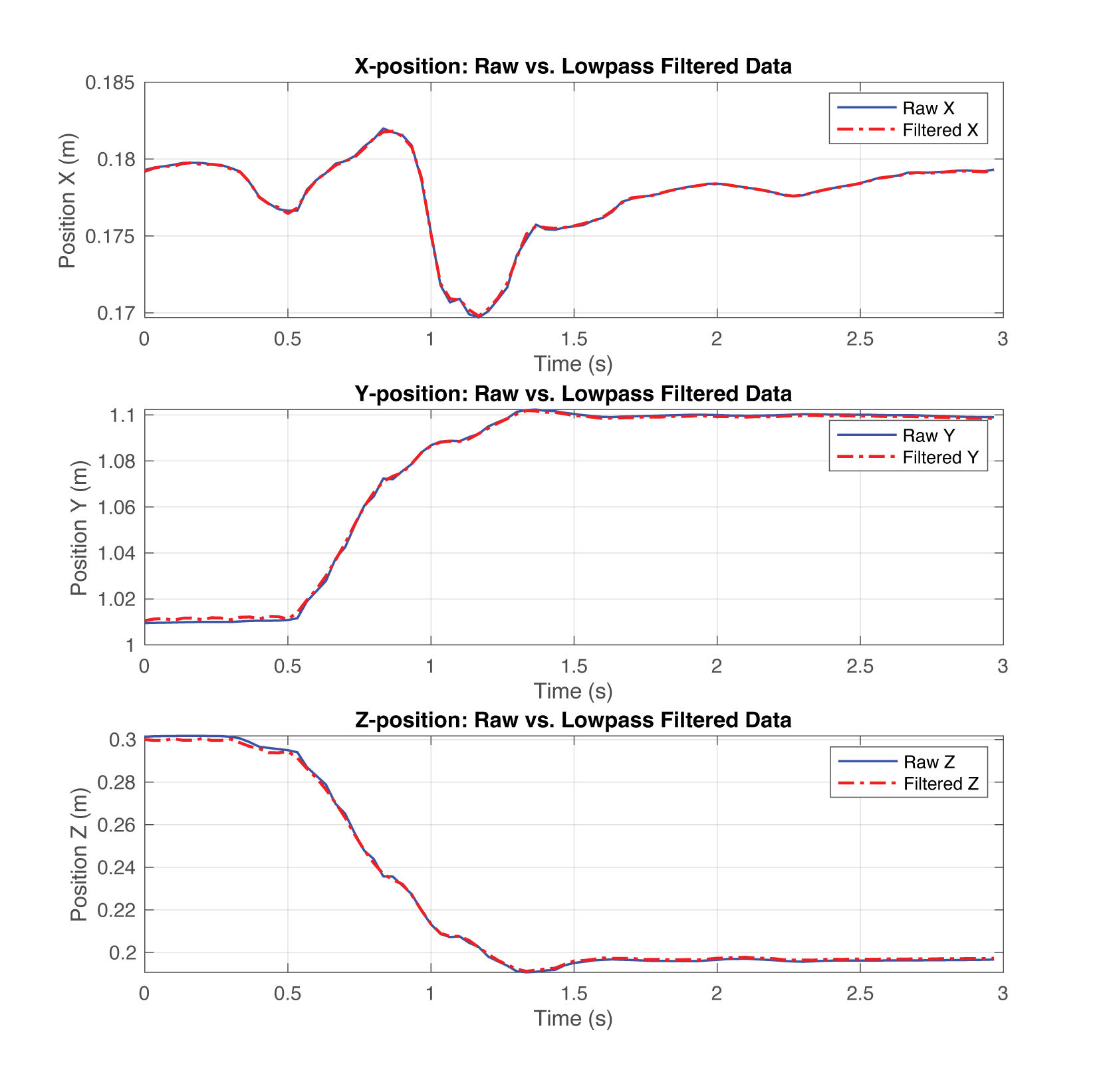

***Figure S1.*** *Raw and low-pass filtered hand position data (Control 7, Target 1, Trial 1). Hand position data in the x-, y-, and z-directions were low-pass filtered at 10 Hz using MATLAB’s ‘lowpass’ function to reduce potential noise. Example data are shown for one representative trial.*
